# Beta HPV8 E6 Induces Micronuclei Formation and Promotes Chromothripsis

**DOI:** 10.1101/2022.02.03.479074

**Authors:** Dalton Dacus, Steven Stancic, Sarah R. Pollina, Elizabeth Riforgiate, Rachel Palinski, Nicholas A. Wallace

**Author notes:** Denotes the corresponding author.

## Abstract

Cutaneous beta genus human papillomaviruses (β-HPV) are suspected to promote the development of non-melanoma skin cancer (NMSC) by destabilizing the host genome. Multiple studies have established the genome destabilizing capacities of β-HPV proteins E6 and E7 as a co-factor with UV. However, the E6 protein from β-HPV8 (HPV8 E6) induces tumors in mice without UV exposure. Here, we examined a UV-independent mechanism of HPV8 E6-induced genome destabilization. We showed that HPV8 E6 reduced the abundance of anaphase bridge resolving helicase, Bloom syndrome protein (BLM). The diminished BLM was associated with increased segregation errors and micronuclei. These HPV8 E6-induced micronuclei had disordered micronuclear envelopes yet retained replication and transcription competence. HPV8 E6 decreased antiproliferative responses to micronuclei and time-lapse imaging revealed HPV8 E6 promoted cells with micronuclei to complete mitosis. Finally, whole genome sequencing revealed that HPV8 E6 induced chromothripsis in 9 chromosomes. These data provide insight into mechanisms by which HPV8 E6-induces genome instability independent of UV exposure.

**Importance:** Some beta genus human papillomaviruses (β-HPVs) may promote skin carcinogenesis by inducing mutations in the host genome. Supporting this, the E6 protein from β-HPV8 (8E6) promotes skin cancer in mice with or without UV exposure. Many mechanisms by which 8E6 increases mutations caused by UV have been elucidated, but less is known about how 8E6 induces mutations without UV. We address that knowledge gap by showing 8E6 causes mutations stemming from mitotic errors. Specifically, 8E6 reduces the abundance of BLM, a helicase that resolves and prevents anaphase bridges. This hinders anaphase bridge resolution and increases their frequency. 8E6 makes the micronuclei that can result from anaphase bridges more common. These micronuclei often have disrupted envelopes yet retain localization of nuclear-trafficked proteins. 8E6 promotes the growth of cells with micronuclei and causes chromothripsis, a mutagenic process where hundreds to thousands of mutations occur in a chromosome.

## Introduction

Non-melanoma skin cancer (NMSC) is the most prevalent type of cancer and more occurrences are noted each year (1–4). While not often deadly, the cost of treating NMSC is $4.8 billion annually in the United States alone (5). Environmental factors play an outsized role in NMSC development, with UV exposure acknowledged as the main etiological agent (6). Age and immunosuppression are additional contributors (7, 8). An increase in risk associated with immunosuppression indicates that an infectious agent can promote NMSC(9). Among possible infectious agents, some members of the beta genus human papillomaviruses (β-HPV) are considered likely candidates because they promote NMSC development in people with a genetic disorder (epidermodysplasia verruciformis) or patients undergoing immunosuppressive therapy (10–12). However, the extent that β-HPV infections contribute to NMSC development in the general population is unclear.

The association between β-HPV infections and NMSC is supported by *in vitro* systems, epidemiological studies, and animal models (13–15). These data were most compelling for a subset of β-HPVs (HPV5, HPV8, and HPV38). They also demonstrated that β-HPV infections rarely persist. However, β-HPV viral loads peak early in NMSC development before becoming dramatically reduced in NMSC cells (16). These observations caused speculation that β-HPV infections introduce tumorigenic mutations capable of independently driving tumorigenesis without continued viral gene expression (17, 18). *In vitro* and *in vivo* studies have shown that the E6 protein from HPV5, HPV8, and HPV38 hinder DNA repair after UV exposure, suggesting the hypothesized role of β-HPV in NMSC is feasible (19, 20). Other studies have indicated a broader role for β-HPV proteins by showing HPV8 E6 (8 E6) causes tumors in mice without UV exposure (21–25). These studies suggest that 8 E6 can introduce genome destabilization without an exogenous stimulus.

Genome integrity is also at-risk during mitosis, as cellular DNA must be replicated and segregated faithfully to avoid mutations. We have shown that 8 E6 attenuates cellular responses that protect genome integrity during mitosis and increases the frequency of aneuploidy (26–28). Aneuploidy results from the unequal segregation of chromosomes suggest that 8 E6 may cause other types of genome instability caused by segregation errors. Micronuclei are one type of destabilization associated with segregation errors. These small enveloped extra-nuclear structures can form when a chromosome gets caught between the nuclei of forming daughter cells in a structure known as an anaphase bridge (29–31). Mutations in micronuclear DNA are common because micronuclear envelopes are unstable limiting the ability to localize cellular factors necessary for replication, transcription, nuclear protein localization, and DNA repair (32–34). It is further mutagenic for a cell to enter mitosis with unrepaired micronuclear DNA. When this occurs, the micronuclear DNA is prone to undergo chromothripsis, where hundreds to thousands of clustered mutations occur over a single cell cycle (35–39). If 8 E6 promotes chromothripsis, it would provide a plausible mechanism by which β-HPV infections as brief as one cell cycle could cause enough mutations to drive NMSC development. Notably, chromothripsis-like events have been identified in NMSCs although there was no attempt to determine if they were caused by a β-HPV infection (40, 41).

Here, we show that 8 E6 hinders chromosome segregation causing an increase in anaphase bridge formation and impeding their resolution. We link this increase to a reduction of Bloom syndrome protein (BLM) and a reduced ability to localize to anaphase bridges. BLM helicase prevents anaphase bridge formation and is critical for their resolution (42, 43). Mechanistically, we provide evidence that the reduction in BLM is the result of reduced ATR-Chk1 kinase activity that stabilizes BLM by blocking cullin-3-mediated degradation (44). These data expand the significance of previous work demonstrating that 8 E6 reduces ATR-Chk1 activation (45–47). Although only a partial decrease in BLM abundance, we identified a significant impact from the BLM reduction as 8 E6 increases the frequency of segregation errors and micronuclei. We go on to characterize these micronuclei, showing that they are more likely to have disordered micronuclear envelopes and that 8 E6 makes it less likely that a cell undergoes antiproliferative responses to micronuclei (i.e., p53 accumulation, caspase activation, and senescence). 8 E6 also increases the frequency that cells with micronuclei undergo mitosis. Finally, we use whole-genome sequencing to demonstrate that 8 E6 causes chromothripsis.

## Results

### p300-dependent BLM reduction correlates with loss of ATR and Chk1 abundance

Because HPV8 E6 (8 E6) reduces activation of the ATR-Chk1 signaling pathway that facilitates BLM stabilization, we hypothesized that 8 E6 decreased BLM abundance (45–47). To evaluate this, we compared BLM abundance in vector control (iHFK LXSN) and HA-tagged 8 E6 expressing (iHFK 8 E6) hTERT-immortalized keratinocytes cells. 8 E6 expression was confirmed by probing for the HA-tag (Supplementary Figure 1A). As expected, 8 E6 reduced ATR-Chk1 signaling (Figure 1A). Consistent with our hypothesis, 8 E6 also decreased BLM abundance. To determine if the reduced BLM required TERT immortalization, we repeated this experiment in primary foreskin keratinocyte cells (HFK LXSN and HFK 8 E6). Expression of 8 E6 in these cells was previously confirmed (27). To further confirm 8 E6 expression, we detected p300 destabilization by immunoblot. (Supplementary Figure 1B). Examination of these cell lines demonstrated that the reduction in BLM levels did not require TERT immortalization (Figure 1B). Further, these cell lines were from different donors indicating that the phenotype is conserved in at least two genetic backgrounds. We take this approach of dual confirmation throughout this manuscript with the goal of similarly probing the requirement of TERT immortalization and the conservation of phenotypes between donors.

**Figure 1.**
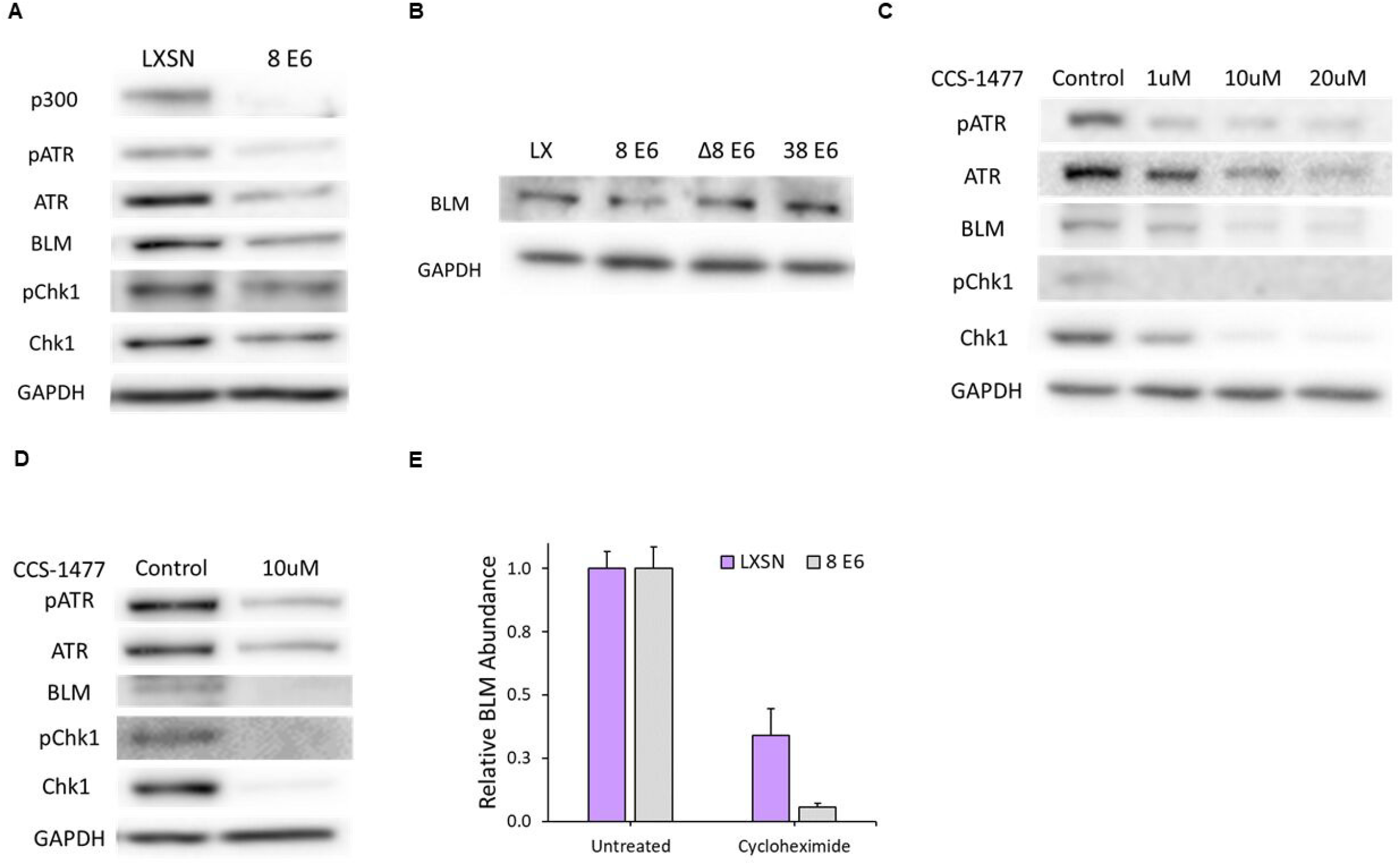
HPV8 E6 reduces BLM abundance. (A) Immunoblots of whole cell lysates from iHFK cells expressing LXSN and 8 E6. (B) Immunoblots of whole cell lysates HFK cells expressing LXSN and the indicated E6 genes (C) Immunoblots of whole cell lysates from iHFK LXSN cells treated with increasing amounts of CCS-1477 for 24 hrs. (D) Immunoblot of whole cell lysates from HFK LXSN cells following 24 hr CCS-1477 treatment. (E) Densitometry of in iHFK LXSN and iHFK 8 E6 cells treated 50 μg/ml cycloheximide for 30 hrs. The graph depicts the mean ± standard error of the mean from three independent experiments. Immunoblots are representative of at least three independent experiments. In Δ8 E6, residues 132 to 136 were deleted from 8 E6.

8 E6 disrupts ATR-Chk1 signaling by binding and destabilizing p300 (47), leading us to hypothesize that 8 E6 decreased BLM levels by binding p300. To test this hypothesis, we determined whether the residues responsible for p300 binding were required for 8E6 to reduce BLM abundance by expressing a mutant 8 E6 without the p300 binding site (HFK 8 E6 Δ132-136) in primary human foreskin keratinocytes. Expression of this mutated 8 E6 in these cells was validated previously, but could not be confirmed by immunoblotting for p300 as HFK 8 E6 Δ132-136 does not degrade p300 (27). BLM levels were not significantly reduced in HFK 8 E6 Δ132-136 cells compared to HFK LXSN cells. To further probe the role of p300 binding in BLM reduction, we expressed HPV38 E6 in primary foreskin keratinocytes (HFK 38 E6). Expression of 38 E6 in these cells has also been confirmed and is not amenable to indirect validation by detecting p300 as 38 E6 weakly binds p300 and does not destabilize p300 (27, 48). Consistent with the requirement for robust binding a p300, 38 E6 also did not reduce BLM abundance (Figure 1B). Deletion of the p300 binding domain disrupts other functions of 8 E6 (23), so we verified the requirement of p300 for optimal BLM abundance by treating iHFK LXSN cells with a small molecule inhibitor of p300 (CCS-1477; (49)). CCS-1477 reduced BLM and ATR-CHK1 activation in a dose-dependent manner (Figure 1C; (47)). CCS-1477 similarly reduced BLM abundance and ATR-Chk1 signaling in HFK cells (Figure 1D). Because Chk1 increases BLM abundance by stabilizing the protein, we hypothesized that 8 E6 decreased BLM stability. To test this, we determine the impact of cycloheximide (protein synthesis inhibitor) on BLM abundance in iHFK LXSN and iHFK 8 E6 cells. Cycloheximide resulted in a greater reduction in BLM abundance in iHFK 8 E6 cells compared to iHFK LXSN cells, consistent with our hypothesis (Figure 1E). The difference, however, did not reach statistical significance (*p*=0.11).

### 8 E6 reduces BLM at anaphase bridges increasing unresolved bridge frequency

Because BLM is required for faithful chromosome segregation (42–44), we hypothesized that 8 E6 increased the frequency of segregation errors. To test this, iHFK cells were synchronized by thymidine block and released to enrich for cells in anaphase. Then, we used immunofluorescent microscopy to identify anaphase cells and determine if they had segregation errors. When identified, segregation errors were grouped into established categories. CENP-A, a centromere marker, was used to distinguish between acentric and lagging chromosomes (Figure 2A). 8 E6 increased the frequency of segregation errors in general and, more specifically, anaphase bridges (Figure 2B). To determine if 8 E6 reduced the amount of BLM that localized to these bridges, we compared BLM staining intensity at chromatin bridges between iHFK LXSN and iHFK 8 E6 cells (Figure 2C). 8 E6 decreased BLM staining at chromatin bridges (Figure 2D). Because BLM is required for anaphase bridge resolution, we hypothesized that 8 E6 increased the time it takes to resolve anaphase bridges. To test this, we compared anaphase bridge duration by time-lapse imaging between iHFK LXSN and iHFK 8 E6 cells expressing a mCherry-tagged H2B for DNA visualization (Movie S1, S2, and Supplementary Figure 2A). Anaphase bridges were more persistent in iHFK 8 E6 cells with one bridge lasting the duration of the observed 30 hours (Figure 2E). However, difficulty in observing these rare events by time-lapse imaging prevented us from collecting enough data to establish statistical significance (*p*=0.36).

**Figure 2.**
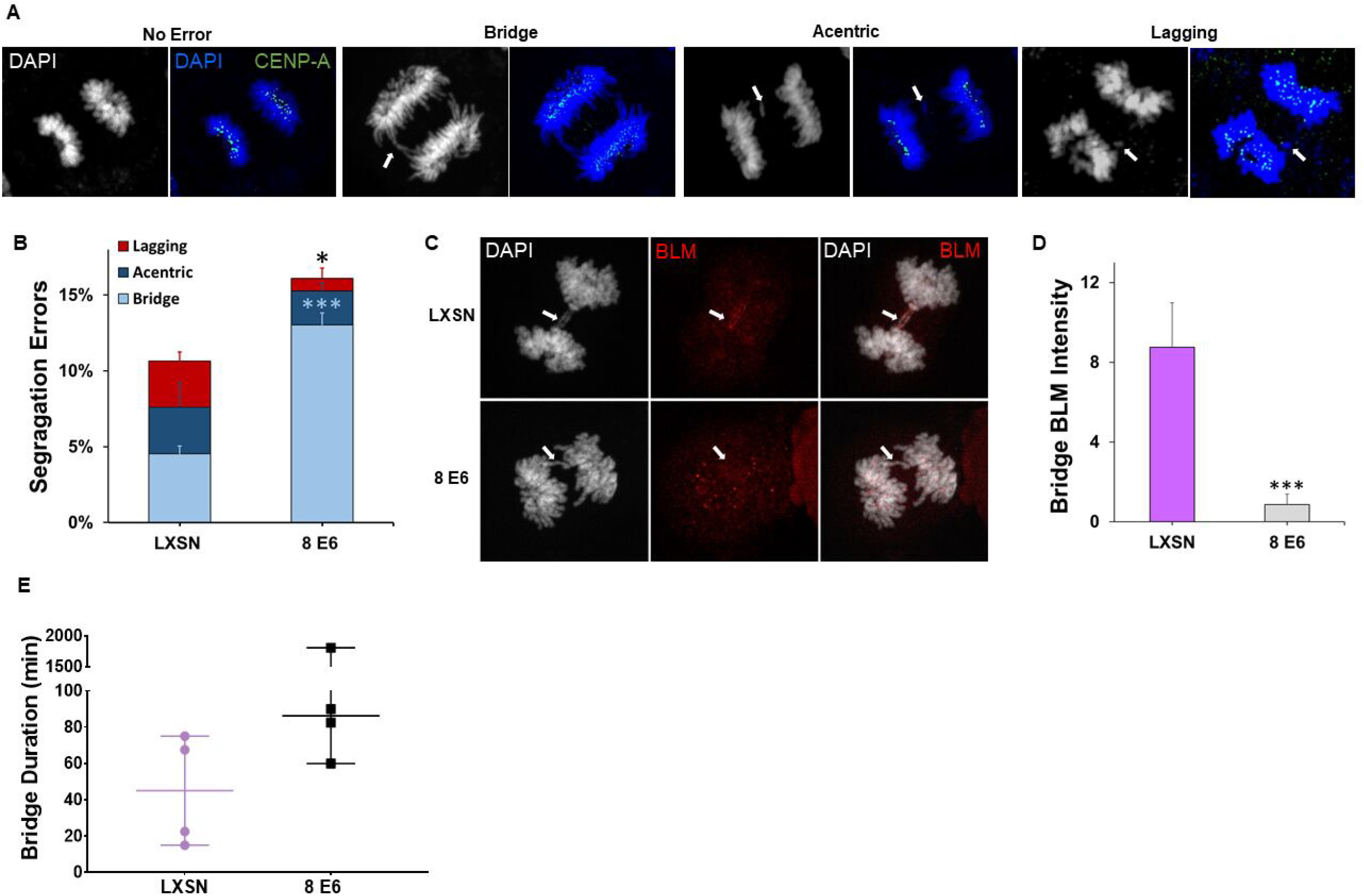
HPV8 E6 augments segregation error rate independent of p300. (A) Images of iHFK cells displaying anaphases with no error (left), a chromatin bridge (left middle), acentric DNA (right middle), and a lagging chromosome (right) stained with DAPI (white and blue) and CENP-A (green); each indicated by arrows. (B) Quantification of segregation errors in iHFK cells after synchronization by thymidine block and released to enrich for mitotic cells. n = at least 221 micronuclei from three experiments. (C) Representative images of BLM (red) localization to chromatin bridges stained with DAPI (white) in iHFK LXSN and 8 E6 cells. Arrows indicate the location of anaphase bridge (D) Mean intensity of BLM staining at chromatin bridges in iHFK LXSN and 8 E6 cells after being synchronized by thymidine block and released to enrich for mitotic cells. n = at least 39 bridges from three experiments. (E) Quantification of time-lapse anaphase bridge duration in iHFK cells. n =4 across three independent experiments. Graphs (B&D) depict mean ± standard error of the mean from three independent experiments except. Graph E depicts the median with a 95% confidence interval from three independent experiments. Asterisks denote significant differences relative to LXSN. * = *p*≤0.05 and *** = *p*≤ 0.001 (Student’s *t*-test).

### 8 E6 increases micronuclei frequency, in part through BLM reduction

Because segregation errors associated with reduced BLM availability can result in the formation of micronuclei (37, 50–53), we hypothesized that 8 E6 expression increased the frequency of micronuclei. To test this, micronuclei were identified using immunofluorescence microscopy (Figure 3A). Micronuclei were more common in both HFK 8 E6 and iHFK 8 E6 cells compared with their LXSN counterparts (Figure 3B and 3C). To provide mechanistic insight, we examined the p300 dependence of the increase in micronuclei. Compared to HFK LXSN cells, both HFK 8 E6 Δ132-136 and HFK 38 E6 cells had a higher frequency of micronuclei. However, micronuclei were significantly more common in HFK 8 E6 compared to HFK 8 E6 Δ132-136 cells.

**Figure 3.**
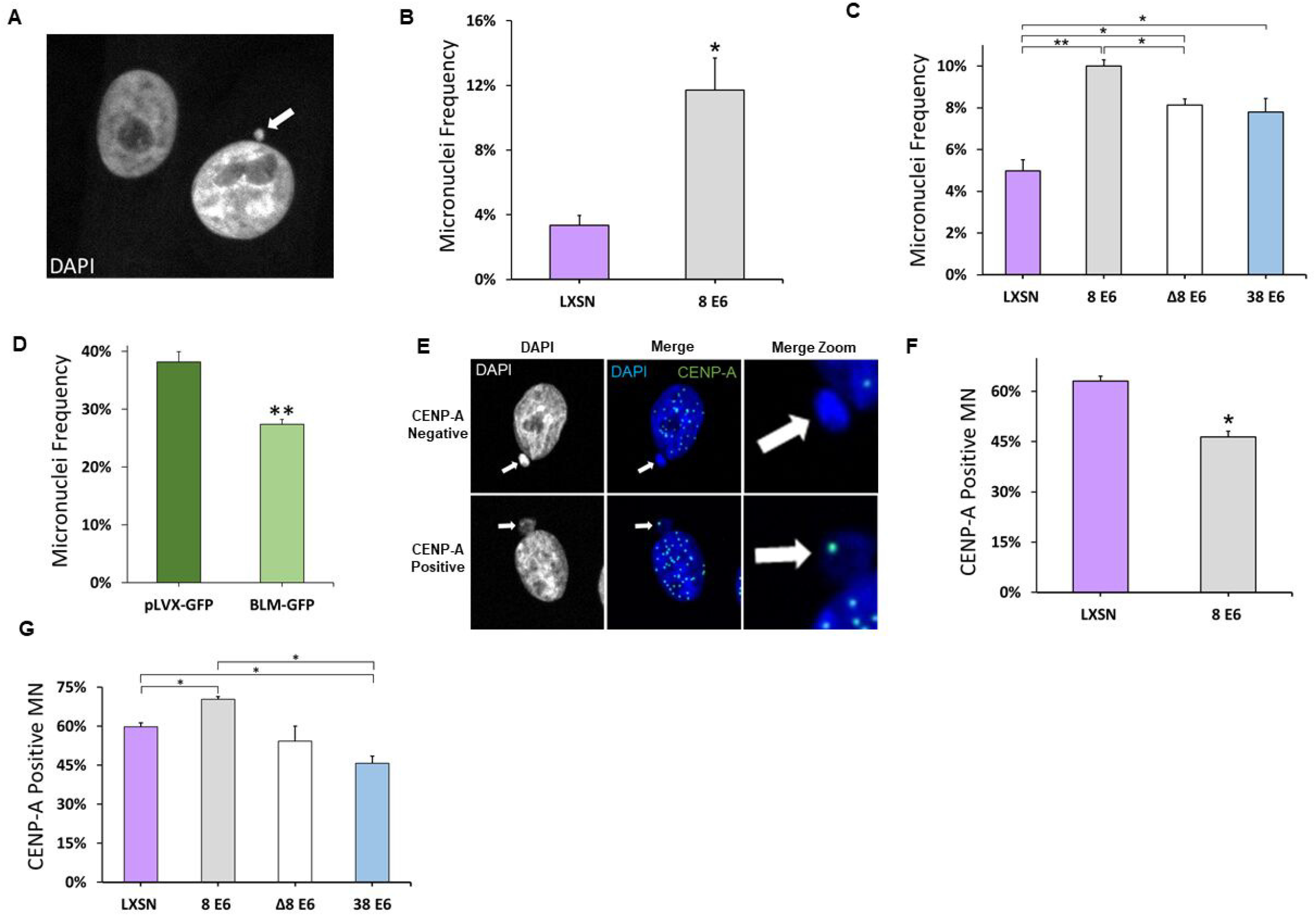
HPV8 E6 increases micronuclei frequency. (A) Representative images of DAPI (white) stained HFK cells, one with (indicated by arrow) and one without a micronucleus. Micronuclei frequency of (B) iHFK and (C) HFK cells. (D) Micronuclei frequency of U2OS 8 E6 cells 72 hours post-transfection with BLM-GFP and pLVX-GFP. >150 cells/cell line were quantified for micronuclei frequency across three independent experiments. (E) Representative images of CENP-A-positive and -negative micronuclei (indicated by arrows) stained for DAPI (white and blue) and CENP-A (green). Percentage of micronuclei with at least one CENP-A foci in (F) iHFK and (G) HFK LXSN and β-HPV E6 cells. At least 250 micronuclei/cell line were quantified for CENP-A frequency across three independent experiments. Graphs depict mean ± standard error of the mean; *n*≥ 3. Asterisks denote significant difference relative to LXSN unless specified with a bar. * = *p*≤ 0.05 and ** = *p*≤0.01 (Student’s *t*-test). In Δ8 E6, residues 132 to 136 were deleted from 8 E6.

To determine the extent that reduced BLM abundance contributed to the elevated frequency of micronuclei, we exogenously expressed a GFP-tagged BLM or GFP alone (transfection control) in U2OS 8 E6 cells. We were unable to test the BLM-dependence of 8 E6-induced micronuclei formation in HFK or iHFK cells as these cell lines were not amenable to BLM transfection. We also confirmed the ability of 8 E6 to increase the prevalence of micronuclei in these cells (Supplementary Figure 3A). Cells were fixed 72 hours post-transfection and DAPI stained to detect micronuclei. This allowed the exogenous BLM to be expressed as cells pass through mitosis 1 to 3 times. We only examined cells expressing GFP to ensure that only transfected cells were considered. Exogenous GFP-BLM expression reduced the frequency of micronuclei in U2OS 8 E6 cells (Figure 3D).

Micronuclei contain either whole (centric) or partial (acentric) chromosomes depending on how they form (54–57). Unrepaired DSBs lead to smaller acentric micronuclei (58), whereas segregation errors result in larger centric micronuclei (59, 60). Our group and others have shown that 8 E6 attenuates DSB repair (19). Here, we demonstrated that 8 E6 can cause segregation errors (Figure 2B). Therefore, we hypothesized 8 E6 caused both acentric and centric micronuclei. To test this we probed for CENP-A, a widely used marker for centric micronuclei, by immunofluorescence microscopy (61). 8 E6 caused acentric and centric micronuclei in iHFK and HFK cells (Figure 3E-3G). Micronuclei produced by 8 E6 were on average significantly smaller than vector control (Supplementary Figure 4A). This, along with 8 E6 micronuclei containing a DSB marker (γH2AX) are consistent with some of these micronuclei forming because of impaired DSB repair (Supplementary Figure 4B and 4C).

### 8 E6 reduces Lamin B1 integrity of micronuclei

Anaphase bridges tend to produce micronuclei lacking micronuclear envelope integrity (62). Therefore, we hypothesized that 8 E6-induced micronuclei would more frequently have unstable micronuclear envelopes. To evaluate micronuclear envelope integrity, we used immunofluorescent microscopy to group micronuclei based on Lamin B1 morphology into established categories (32). First, micronuclei were placed into two large groups based on whether Lamin B1 staining displayed clear peripheral localization around the micronucleus (intact) or not (disordered). Micronuclei with intact membranes were further segregated based on whether they had a continuous (no holes) or discontinuous (one or more holes) nuclear Lamin B1 staining (Figure 4A; top two panels). Micronuclei with disordered micronuclear membranes were also further split based on whether nuclear Lamin B1 staining appeared collapsed or was absent (Figure 4A; bottom two panels). The ratio of intact to disordered micronuclear membranes in iHFK LXSN cells were similar to prior reports in other cell lines (Figure 4B) (32, 63). In contrast, micronuclear membranes in iHFK 8 E6 cells were more often disordered, with a specific increase in micronuclei without a membrane (Figure 4B). Disordered membranes typically prevent nuclear factors from being imported from the cytoplasm (32, 51, 64). To characterize the impact of micronuclear membrane integrity on the localization of nuclear proteins, we generated iHFK LXSN and iHFK 8 E6 cells that stably expressed a fusion gene consisting of two RFP proteins and a nuclear localization signal (RFP-NLS). We examined Lamin B1 morphology and RFP staining in these cells and found that RFP localized to micronuclei with intact envelopes (Figure 4C). 8 E6 increased the proportion of RFP-positive micronuclei with discontinuous envelopes. This suggests 8 E6 disrupts envelope integrity but retains loss of nuclear localization when the envelope becomes disordered.

**Figure 4.**
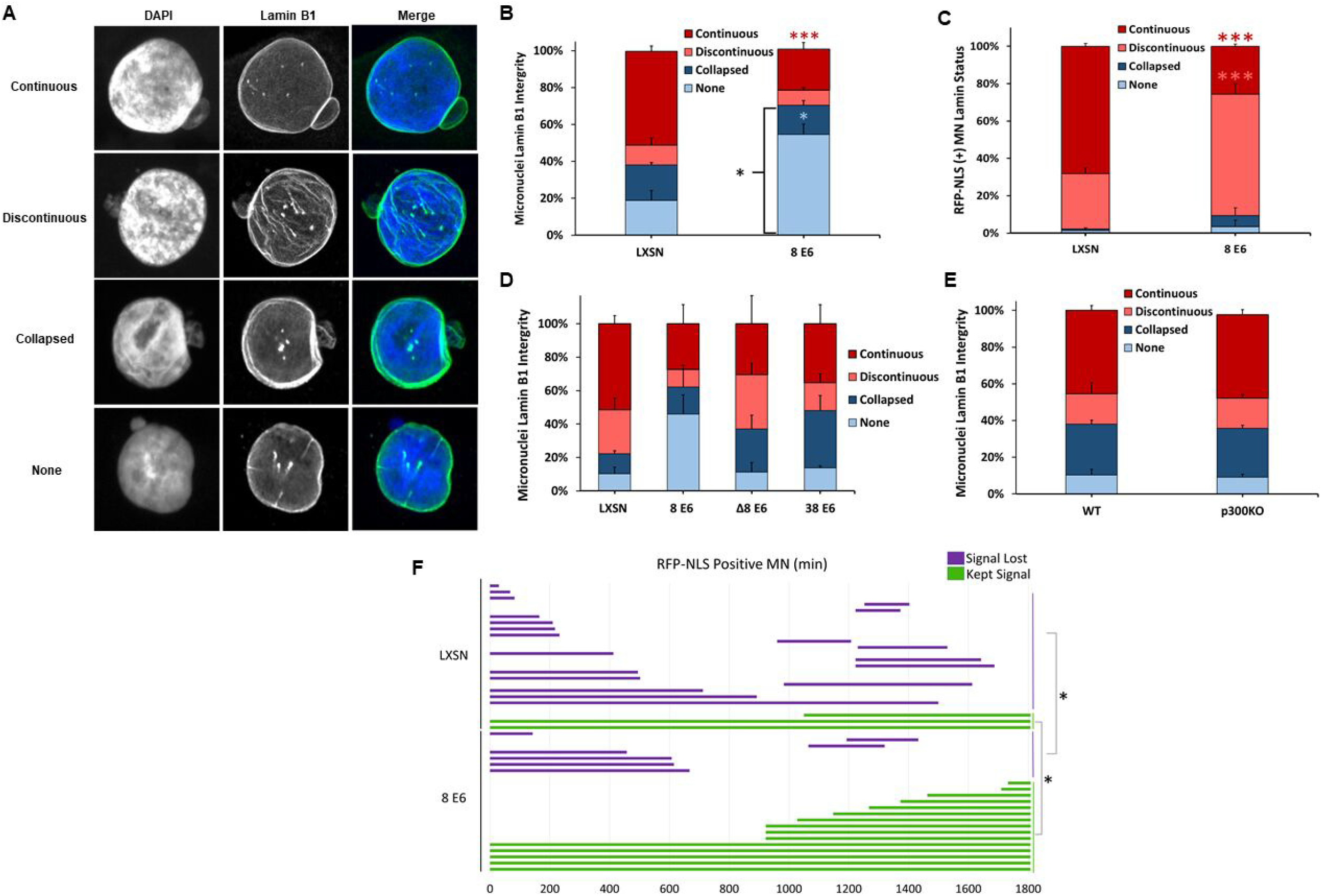
HPV8 E6 reduces Lamin B1 integrity of micronuclei. (A) Representative images of cells with intact (continuous and discontinuous) and disordered (collapsed and none) micronuclear envelopes. DNA is stained with DAPI (blue) and nuclear membranes were detected with antibodies against Lamin B1 (green). (B) Quantification of Lamin B1 micronuclear morphology in asynchronous iHFK cells. n ≥ 234 micronuclei from three independent experiments. (C) Quantification of RFP-NLS-positive micronuclei segregated by micronuclear morphology (as determined by Lamin B1 staining) in iHFK cells. (D) Quantification of Lamin B1 micronuclear morphology in asynchronous HFK cells. n = at least 146 micronuclei from three experiments. Same as (B) but in HCT 116 cells. Quantification of Lamin B1 micronuclear morphology in asynchronous iHFK cells. n ≥ 200 micronuclei from three independent experiments. (F) Length of time (min) in which RFP-NLS-positive micronuclei in iHFK RFP-NLS cells persisted for 30 hours. Red, salmon, and blue asterisks denote a significant difference from LXSN in respect to continuous, discontinuous, and none Lamin B1 morphologies, respectively. Graphs depict the mean ± standard error of the mean from three independent experiments. Asterisks denote significant differences relative to LXSN. * = *p*≤0.05 and *** = *p*≤ 0.001 (Student’s *t*-test). In Δ8 E6, residues 132 to 136 were deleted from 8 E6.

To determine if 8 E6 could reduce micronuclear membrane stability in HFK cells, we examined micronuclear membranes by immunofluorescence microscopy. While not statistically significant (p=0.079), these data were similar to the observations in iHFK cells (Figure 4D). To determine the extent that p300 destabilization contributed to this phenotype, we compared membrane integrity between HFK 8 E6, HFK 8 E6 Δ132-136, and HFK 38 E6 cells, but they did not demonstrate a role for p300. The frequency of micronuclei without membranes in these cells were like the frequency in HFK LXSN (Figure 4D). However, the prevalence of collapsed micronuclear membranes was elevated in HFK 8 E6 Δ132-136, and HFK 38 E6 cells. These differences did not reach statistical significance. We also did not find a difference in micronuclear envelope integrity when comparing HCT 116 cells with or without p300 (Figure 4E). Thus, we were unable to draw a conclusion about the role of p300 destabilization in 8 E6-mediated disruption of micronuclear membranes.

Notably, 8 E6 increased the frequency of RFP-NLS-positive micronuclei with discontinuous envelopes, but there was negligible RFP in micronuclei with disordered envelops (Figure 4C). Because discontinuous micronuclear envelopes are prone to become disordered with time (25), we hypothesized that 8 E6 increased the chance that cells lost nuclear RFP localization over time. To test this, we used time-lapse imaging to track RFP-NLS status of micronuclei (retained or lost) for 30 hours in iHFK LXSN and 8 E6 cells (Movie S3 and S4). Instead, we found that 8 E6 increased the retention of nuclear localization (Figure 4F). Together, these data indicate that 8 E6 promotes nuclear localization despite more frequently having damaged micronuclear envelopes.

### 8 E6 promotes some nuclear function despite disrupted micronuclear envelopes

Membrane integrity is critical for the localization of proteins involved in nuclear functions (replication, transcription, and DNA damage repair). Failure to localize nuclear factors to micronuclei increases the likelihood that micronuclei contain damaged DNA (32, 51, 65). We used immunofluorescence microscopy to characterize these processes in the micronuclei of HFK LXSN and HFK 8 E6 cells. We first evaluated active transcription and replication in these micronuclei using histone H3 acetylated at K27 (H3K27ac) and BrdU incorporation as respective markers. Consistent with previous results (32), transcription and replication were reduced in micronuclei with disordered membranes in HFK LXSN cells (Figure 5A and 5B, and Supplementary Figure 5A and 5B). 8 E6 did not cause robust changes in transcription or replication in micronuclei with intact membranes. However, 8 E6 allowed both replication and transcription to occur significantly more frequently in micronuclei without envelopes. To evaluate the DDR in micronuclei, we exposed cells to a radiomimetic (zeocin) and assessed the formation of DSB repair foci (53bp1). Consistent with previous reports (32, 33) we found the DDR response is attenuated in micronuclei with disordered envelopes in HFK LXSN and HFK 8 E6 cells (Figure 5C and Supplementary Figure 5C).

**Figure 5.**
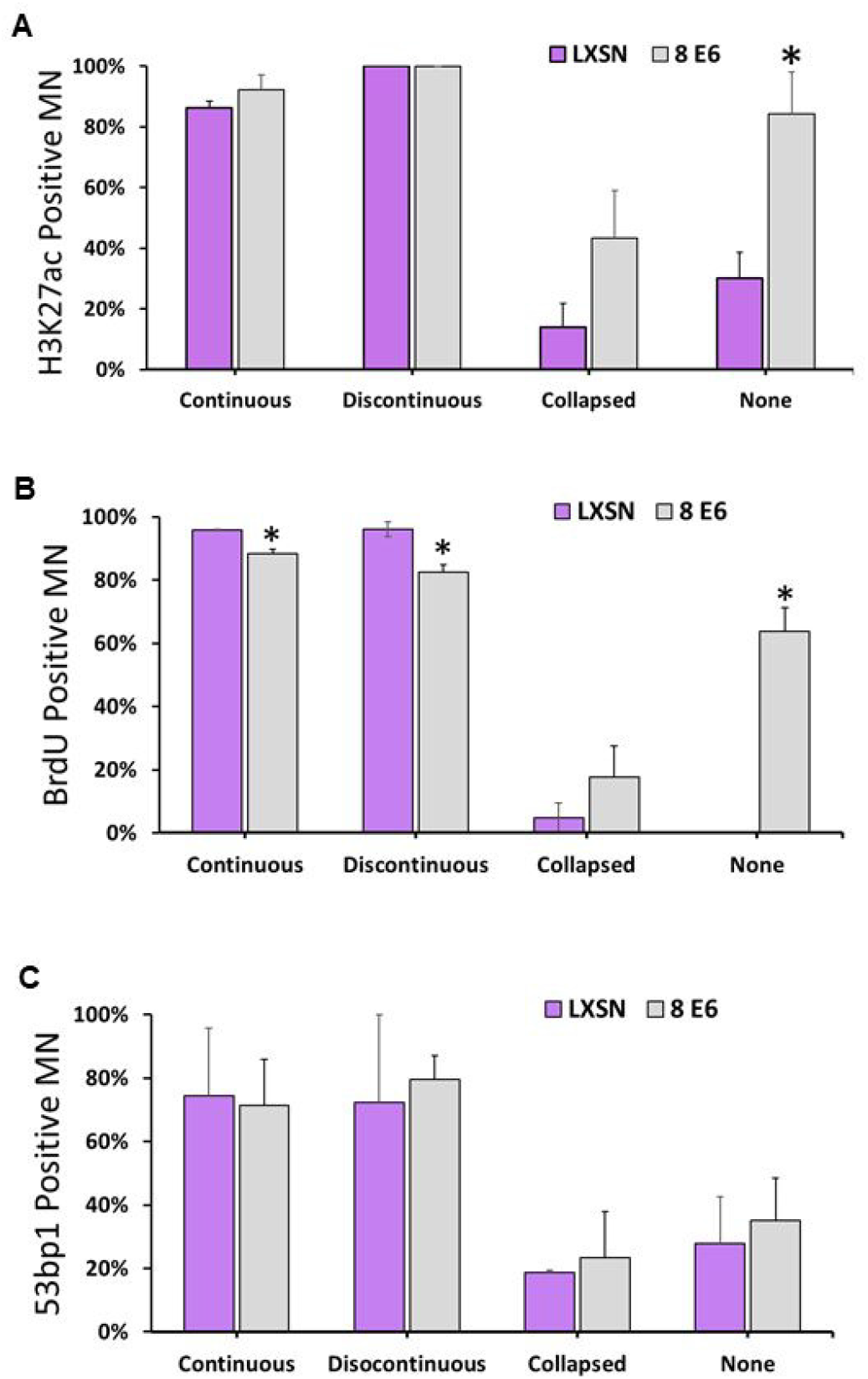
Some micronuclear functions changed by HPV8 E6. Frequency of iHFK cells with (A) acetylated H3K27-positive, (B) 53bp1-positive, and (C) BrdU-positive micronuclei. At least 150 micronuclei/cell line were quantified across three independent experiments. Graphs depict mean ± standard error of the mean; Asterisks denotes significant difference relative to LXSN. * = *p*≤0.05 (Student’s *t*-test). Micronuclei is abbreviated as MN.

### 8 E6 reduces apoptosis and senescence in response to micronuclei

Micronuclei induce apoptosis and senescence (66, 67). Because HPV replication requires cellular proliferation, we hypothesized that 8 E6 and 38 E6 impeded apoptosis and senescence in response to micronuclei. As p53 can facilitate these responses, we compared p53 staining intensity in HFK LXSN, HFK 8 E6, HFK 8 E6 Δ132-136, and HFK 38 E6 cells that did and did not have micronuclei (Figure 6A). The presence of a micronucleus similarly increased p53 staining intensity in HFK LXSN, HFK 8 E6, and HFK 8 E6 Δ132-136 cells (Figure 6B). HFK 38 E6 cells had significantly increased p53 intensity compared to all other HFK cells.

**Figure 6.**
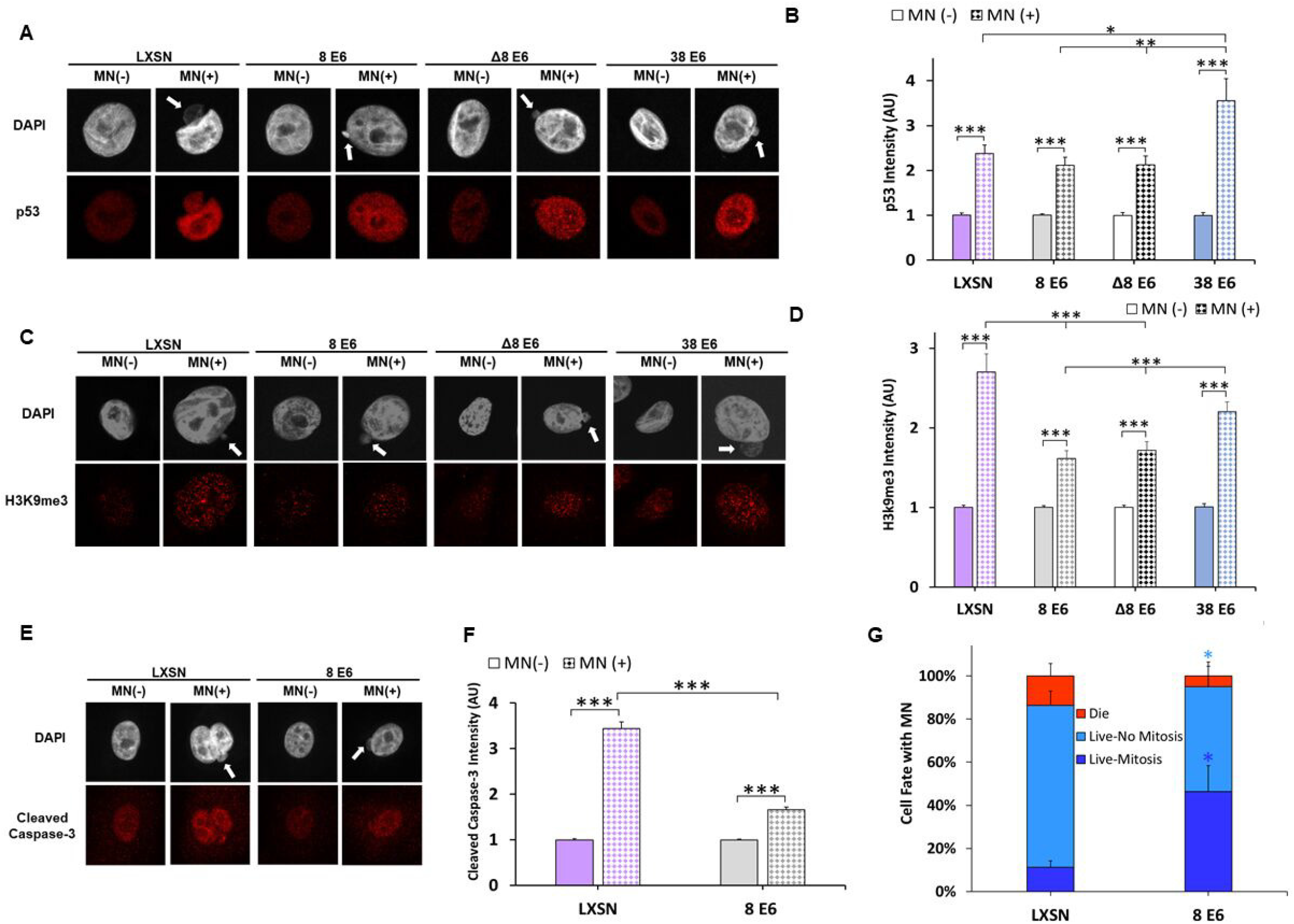
HPV8 E6 reduces a cell’s apoptotic and senescent response to micronuclei. (A) Representative images of HFK cells stained for p53 (red) and DNA (DAPI, white) with (indicated by white arrow) and without a micronucleus. (B) Quantification of normalized p53 intensity in HFK cells with and without micronuclei. (C) Representative images of methylated H3K9-(red) and DAPI (white)-stained HFK cells with (indicated by white arrow) and without a micronucleus. (D) Quantification of normalized methylated H3k9 intensity in HFK cells with and without micronuclei. At least 200 cells/cell line were quantified across three independent experiments. (E) Representative images of HFK cells stained for cleaved caspase-3- (red) and DAP (white) with (indicated by white arrow) and without a micronucleus. (F) Quantification of normalized cleaved caspase 3 intensity in iHFK cells with and without micronuclei. ≥550 cells/cell line were quantified across 4 independent experiments. (G) Quantification of cell fate imaging micronuclei-positive iHFK LaminB1-GFP H2B-RFP cells for 30 hrs. Light and dark blue asterisks denote a significant difference between ‘Live-no mitosis’ and ‘Live-mitosis’ to LXSN, respectively. Graphs depict mean ± standard error of the mean; Asterisks denotes significant difference relative to LXSN unless specified with a bar. * = *p*≤0.05 and ** = *p*≤0.01 (Student’s *t*-test). In Δ8 E6, residues 132 to 136 were deleted from 8 E6. Micronuclei is abbreviated as MN

Next, we used similar approaches to examine markers of senescence (histone H3 methylated at K9; H3K9me3) and apoptosis (cleaved caspase-3). H3K9me3 staining intensity was increased by micronuclei in all cell lines (Figure 6D). However, this increase was significantly reduced in HFK 8 E6 and HFK 8 E6 Δ132-136 cells compared to HFK LXSN cells (Figure 6C-D). H3K9me3 staining in HFK 38 E6 cells more closely resembled the intensity found in HFK LXSN cells. Cleaved caspase-3 staining was likewise elevated in all iHFK cell lines with micronuclei (Figure 6E-F). However, 8 E6 significantly attenuated the increase in cleaved caspase-3 staining in cells with micronuclei (Figure 6E-F). Because 8 E6 attenuates the accumulation of apoptosis and senescent markers in cells with micronuclei, we hypothesized that 8 E6 increased the frequency that cells with micronuclei underwent mitosis. To test this, we transduced LXSN and 8E6 iHFK cells with mCherry-tagged histone H2B to visualize DNA and used time-lapse imaging to track the fate of cells with micronuclei over 30 hours. 8 E6 made it significantly more likely that a cell with a micronucleus completed mitosis (Figure 6G and Movie S5-6).

### 8 E6 promotes chromothripsis events

Micronuclear DNA is at risk of chromothripsis if the cell containing them completes mitosis (34, 36, 39). Because 8 E6 increases the frequency of micronuclei and promotes proliferation (Figure 3B and 6G), we hypothesized that 8 E6 caused chromothripsis. To detect chromothripsis, we performed whole genome sequencing on passaged-matched iHFK LXSN and 8 E6 cells. Raw reads were trimmed for quality and analyzed using a standard pipeline, using the vector mutations as a control (Methods, Supplementary Figure 6). Shatterseek software (68) was used to determine chromothripsis by analyzing copy number changes (CN) and structural variant (SV) differences between cell lines. We identified 9 chromosomes with chromothripsis mutations and 7 others with high numbers of structural variants (Figure 7A-B). Significant instability was not found in eight other chromosomes. We found smaller chromosomes were less likely to have instability following mitosis, however, did have an observed decrease in copy number changes and an increase in structural variants per nucleotide (Figure 7C).

**Figure 7.**
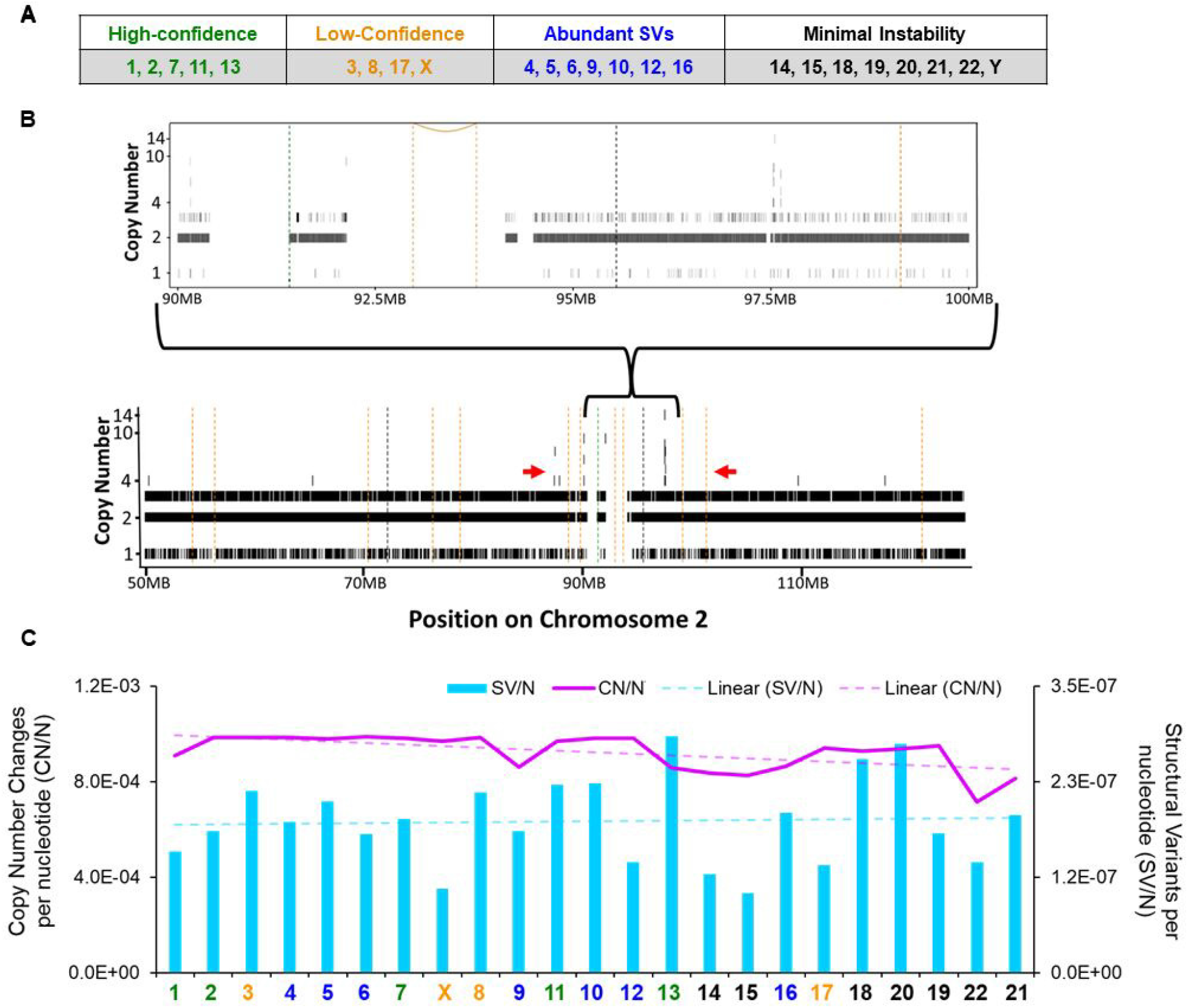
HPV8 E6 promotes non-canonical chromothripsis. (A) Table summarizing the chromothripsis status of iHFK 8 E6 chromosomes analyzed when compared to iHFK LXSN (vector-control) chromosomes. (B) Chromothripsis analysis from chromosome 2. Vertical hashed bars show deletion-like events (orange), head-to-head inversions (black), and tail-to-tail inversions (green). Copy number (CN) variations are indicated by solid black bars ranging from 1-14. The red arrows denote an area with Shatterseek-defined chromothripsis. (C) Total copy number variations per nucleotide (CN/N) and structural variants per nucleotide (SV/N) per chromosome in iHFK 8 E6 cells. The purple line represents CN/N and the blue bars represent SV/N. Chromosome color corresponds to chromothripsis status from (A).

## Discussion

β-HPV research often focuses on the ability of the β-HPV proteins to facilitate the accumulation of UV-induced mutations (69, 70). However, a growing body of evidence has demonstrated that 8 E6 also hinders genome integrity independent of UV exposure (26–28). In this study, we show 8 E6 causes anaphase bridges by reducing BLM abundance. (Figure 1–2). In turn, this leads to an increased frequency of micronuclei (Figure 3). These micronuclei are less likely to have intact membranes, but surprisingly retain the localization of nuclear proteins and processes (Figure 4–5). 8 E6 attenuates antiproliferative responses associated with the micronuclei it generates, allowing cells to complete mitosis (Figure 6). Our whole genome sequencing data suggest these phenotypes culminate in significant genome destabilization in the form of chromothripsis (Figure 7).

Among HPVs, HPV8 is not alone in its ability to induce genomic instability in association with mitotic defects and micronuclei. In fact, this capability spans across two HPV genera (alpha and beta) (71, 72). The E6 oncoprotein from the alpha genus human papillomavirus 16 (HPV16 E6) also promotes micronuclei and anaphase bridges (73, 74). Micronuclei are also associated with alpha HPV oncogene expression in human tissues, where they can distinguish between HPV-positive and HPV-negative tumors (75, 76). HPV16 oncoprotein E7 (HPV16 E7) exacerbates HPV16 E6-induced anaphase bridges and micronuclei (73, 74, 77). Notably, the combined expression of HPV38 E6 and E7 has previously been shown to promote genomic instability associated with increased anaphase bridges (71). Our data show that HPV38 E6 can increase the frequency of micronuclei without E7 (Figure 3C). This is consistent with other published data that demonstrate multiple β-HPV E6s can extensively alter the cellular environment without co-expression of their corresponding E7s (19, 69). While the ability of HPV16 E6/E7 and HPV38 E6/E7 to increase chromosomal instability and anaphase bridge prevalence is impaired by telomerase expression, we show that the ability of 8 E6 to cause similar phenotypes was unaffected by telomerase expression (71, 73). It is unclear whether the differences should be attributed to the lack of 8 E7 in our experimental system or whether they represent differences in 8 E6 and other HPV E6s.

Albeit via differing mechanisms, a common phenotype between HPV16, HPV38 and HPV8 are their capacities to promote cell proliferation despite inhibitory stimuli from the cell (23, 78, 79). This is likely an evolutionarily conserved phenotype due to the requirement that host cells remain proliferatively active for the completion of HPV’s lifecycle. Micronuclei can trigger these inhibitory stimuli, resulting in apoptosis and senescence (80–82). To the best of our knowledge, how HFKs respond to micronuclei had not yet been determined. We show that micronuclei (or the damage associated with them) elicit apoptotic and senescent responses in vector control keratinocytes (Figure 6). HPV8 E6 attenuates cellular responses to micronuclei, promoting entry and completion of mitosis (Figure 6G). This is notable because completion of mitosis greatly increases the risk of chromothripsis (36, 39). If a second mitosis occurs, the chromatin trapped inside the micronucleus often becomes missegregated, resulting in further anaphase bridges and/or micronuclei (83, 84).

The increases in micronuclei and chromothripsis are likely caused by 8 E6-mediated reductions in BLM. BLM mutations cause a predisposition to cancer (85). BLM is an essential genome stabilizer due to its regulation of DNA recombination, replication, and both homologous and non-homologous double-strand break repair pathways (86). We hypothesize that the 8 E6-induced BLM reduction results in yet unreported defects in DNA repair. This could include increased sister chromatid intertwines that result in the anaphase bridges reported here. Another interesting observation from this work is the ability of 8 E6 to promote replication and transcription in micronuclei without envelopes. Whether 8 E6 is actively involved in recruiting nuclear factors or causes this localization indirectly is unclear.

Our whole genome sequencing analysis is the first to directly compare two passage-matched hTERT–immortalized primary cells lines. This is important because we can directly compare and describe the genomic changes following HPV8 E6 exposure. This approach could improve the existing knowledge of HPV16-induced chromothripsis (72) or of mutations that promote chromothripsis. Finally, when compared to previous chromothripsis reports, the frequency of copy number oscillations with 8 E6 expression is higher, and the changed fragment sizes are smaller (68, 87). This observation suggests the use of passage-matched immortalized primary cells lines changes the landscape of chromothripsis calling during analysis and/or tumors caused by HPV8 infection have a distinct mutation pattern.

Further, comparisons between passage-matched non-immortalized primary cell lines are needed to determine the extent that hTERT immortalization is required for 8 E6 to induce chromothripsis. There are other remaining questions as well. Is p300 degradation required for 8 E6 to cause chromothripsis? Do other β-HPV E6s cause chromothripsis? Are the unusual copy number oscillations found in NMSCs?

## Material and Methods

### Cell culture

U2OS and HCT 116 cells were maintained in Dulbecco modified Eagle medium (DMEM) supplemented with 10% fetal bovine serum (FBS) and penicillin-streptomycin. Primary HFK cells were derived from neonatal human foreskins. hTERT-immortalized-HFK (iHFK cells were obtained from Michael Underbrink, University of Texas Medical Branch). HFK and iHFK cells were grown in EpiLife medium supplemented with calcium chloride (60μM), human keratinocyte growth supplement (Thermo Fisher Scientific), or Keratinocyte Growth Medium 2 (PromoCell) supplemented with calcium chloride (60μM), SupplementMix, and penicillin-streptomycin. HPV genes were cloned, transfected, and confirmed as previously described (27).

### Plasmids

2×RFP-NLS (RFP-NLS) was a gift from Emily Hatch and previously described (32) and contains mCherry-TagRFP-NLS in the Gateway vector pDONOR20. GFP-BLM was a gift from Nathan Ellis (Addgene plasmid # 80070; http://n2t.net/addgene:80070; RRID: Addgene_80070) pLV-LaminB1-GFP/H2B-mCherry was purchased from VectorBuilder and is detailed here (https://en.vectorbuilder.com/vector/VB210611-1255jkx.html). (88)

### Immunoblotting

After being washed with ice-cold phosphate-buffered saline (PBS), cells were lysed with radioimmunoprecipitation assay (RIPA) lysis buffer (VWR Life Science) supplemented with phosphatase inhibitor cocktail 2 (Sigma) and protease inhibitor cocktail (Bimake). The Pierce bicinchoninic acid (BCA) protein assay kit (Thermo Scientific) was used to determine protein concentration. Equal protein lysates were run on Novex 4-12% Tris-Glycine WedgeWell mini gels (Invitrogen) and transferred to Immobilon-P membranes (Millipore). Membranes were then probed with the following primary antibodies: glyceraldehyde-3-phosphate dehydrogenase (GAPDH) (Santa Cruz Biotechnologies; catalog no. sc-47724), pATR (Thr1989) (58014S, Cell Signaling Technology), ATR (2790S, Cell Signaling Technology), pCHK1 (Ser345) (133D3) (2348S, Cell Signaling Technology), CHK1 (2G1D5) (2360S, Cell Signaling Technology), p300 (Santa Cruz Biotechnologies; catalog no. sc-584), BLM (2742S, Cell Signaling Technology) and after exposure to the matching horseradish peroxidase (HRP)-conjugated secondary antibody, cells were visualized using SuperSignal West Femto maximum sensitivity substrate (Thermo Scientific).

### Immunofluorescence microscopy

Cells were seeded onto either 96-well glass-bottom plates (Cellvis) or etched-coverslips and grown overnight. Cells were fixed with 4% formaldehyde. Then, 0.1% Triton-X solution in PBS was used to permeabilize the cells, followed by blocking with 3% bovine serum albumin in PBS for 30 min. Cells were then incubated with the following: alpha-tubulin (Abcam; catalog no. ab18251), α-tubulin (3873S, Cell Signaling Technology), p53 (2527S, Cell Signaling Technology), BLM (gift from Matthew Weitzman, Children’s Hospital of Philadelphia), CENP-A (3–19) (GTX13939, GeneTex), Lamin B1 (C-12) (sc-365214), Santa Cruz Biotechnology), phospho-Histone H2AX (Ser139) (20E3) (9718S, Cell Signaling Technology), 53bp1 (4937S, Cell Signaling Technology), Histone H3 (acetyl K27) (ab4729, Abcam), Tri-methyl-Histone H3 (Lys9) (D4W1U) (13969T, Cell Signaling Technology), Cleaved Caspase-3 (Asp175) (9661S, Cell Signaling Technology). The cells were washed and stained with the appropriate secondary antibodies: Alexa Fluor 594 goat anti-rabbit (Thermo Scientific; catalog no. A11012) and Alexa Fluor 488 goat anti-mouse (Thermo Scientific A11001). After washing, the cells were stained with 2μM 4’,6-diamidino-2-phenylindole (DAPI) in PBS and visualized with the Zeiss LSM 770 microscope. Images were analyzed using ImageJ techniques previously described in reference (89).

### Detection of BrdU incorporation

Detection of BrdU was performed the same as immunofluorescent cells with the addition of a 30 min incubation with 1.5 M HCl for 30 min between Triton-X and 3% bovine serum albumin. Primary antibodies as indicated and stained with Alexa Fluor 594 anti-BrdU (BU1/75 (ICR1)) (ab20076, Abcam). After washing, the cells were stained with 2μM 4’,6-diamidino-2-phenylindole (DAPI) in PBS and visualized with the Zeiss LSM 770 microscope. Images were analyzed using ImageJ techniques previously described in reference (89).

### 53bp1 detection

Cells were seeded onto etched coverslips and grown overnight. Cells were exposed to 10 μg/ml of zeocin for 10 min and then washed. 1 hr post zeocin cells were fixed and stained following immunofluorescent microscopy steps above.

### Segregation error

Cells were arrested in G1/S phase by culture in 2mM thymidine for 16 hours, washed to release, and grown for 9 hr 10 min in the absence of thymidine to enhance mitotic cells. Cells were shaken into the media, concentrated, and cytocentrifuged onto glass coverslips. Cells were fixed with 4% formaldehyde. Then, 0.1% Triton-X solution in PBS was used to permeabilize the cells, followed by blocking with 3% bovine serum albumin in PBST supplemented with 300mM Glycine for 1 hr. Primary and secondary antibodies were stained for 16 hr and 1 hr 20 min, respectively. After washing, the cells were stained with 2μM 4’,6-diamidino-2-phenylindole (DAPI) in PBS and visualized with the Zeiss LSM 770 microscope. Images were analyzed using ImageJ techniques previously described in reference (89)

### Establishment of RFP-NLS and LaminB1-GFP/H2B-RFP expressing cells and transient expression of BLM-GFP

RFP-NLS expressing iHFK cells were generated by transfecting with the RFP-NLS expression plasmids using Xfect transfection reagent (Takara Bio; 631317). Cells containing RFP-NLS plasmids were selected using 1 μg/ml puromycin. Drug resistant colonies were pooled after 10 days. LaminB1-GFP/H2B-RFP expressing iHFK cells were generated by transducing with LaminB1-GFP/H2B-RFP lentivirus according to the manufacturer’s protocol then selected with 1 μg/ml puromycin until mock transfected cells ceased proliferation.

### Time-Lapse Imaging

Cells were plated into 6-well glass bottom plates (Cellvis) and imaged on a BioTek LionheartFX Automated Microscope with a 20x air objective at 37°C and 5% CO_2_ for 30 hrs. Images were captured using Gen5 software (Biotek). Videos were cropped and adjusted for brightness and contrast using ImageJ.

### Whole genome mate-pair sequencing

Passage-matched iHFK LXSN and 8 E6 cells were subjected to whole genome sequencing. DNA was extracted from cells using the Qiagen High Molecular Weight (HMW) DNA extraction kit, library prepped using the Illumina DNA prep library prep kit, and sequenced on an Illumina Novaseq using an SP 250PE v1.5 cartridge, by the manufacturer’s specification. The average coverages were 53x and 75x for iHFK LXSN and 8 E6 cells, respectively. The raw reads were analyzed using a validated pipeline with some modifications (87). Most notably mutations in 8 E6 expressing HFK cells were identified by comparison to the sequencing from vector control cells rather than the reference genome (87). Raw reads were trimmed with trimmomatic v0.40, aligned with BWA v0.7.17 to Genbank reference GRCh37 (RefSeq # GCF_000001405.25), and copy numbers (CN) were calculated with Xcavator, using default parameters (90). The alignment was subjected to structural variant (SV) calling in sVABA using a log odds ratio of 4 and quality score above 30 (91). CN and SV variants were used to determine the likelihood of chromothripsis in Shatterseek with default parameters (68). Confidence was called for each chromosome based on the number of coinciding structural and copy number variants, the location of genomic breakpoints, and the frequency of all three parameters coinciding.

### Measurement of protein half-life

iHFK LXSN and 8 E6 cells were grown in 6-well plates until ~80% confluency. Cells were treated with 50 μg/ml cycloheximide for indicated times and whole cell lysates for all time points were harvested together. Western blot analysis same as above.

### Statistical analysis

Unless otherwise noted, statistical significance was determined by an unpaired Student’s t-test and was confirmed when appropriate by two-way analysis of variance (ANOVA) with Turkey’s correction. Only *p* values of less than 0.05 were reported as significant.

## Supplementary Figure Legends

Supplementary Figure 1:

(A) Immunoblots of HA-tagged 8 E6 and corresponding p300 staining in iHFK LXSN and 8 E6 cells. (B) Immunoblot analysis of p300 abundance in HFK cells.

Supplementary Figure 2:

(A) Representative time-lapse images of chromatin bridge resolution in iHFK LXSN and 8 E6 cells transduced with H2B-mCherry (white). Arrows indicate the presence of an anaphase bridge.

Supplementary Figure 3:

(A) Micronuclei frequency of U2OS cells. >136 cells/cell line were quantified for micronuclei frequency across three independent experiments. The graph depicts the mean ± standard error of the mean; Asterisks denotes significant difference relative to LXSN. ** = *p* ≤ 0.01 (Student’s *t*-test).

Supplementary Figure 4:

(A) Percentage of micronuclear area normalized to area of primary nucleus in HFK cells expressing β-HPV E6 proteins. At least 150 micronuclei/cell line were quantified across three independent experiments. (B) Percentage of γH2AX-positive micronuclei in HFK β-HPV E6 cells. At least 75 micronuclei/cell line were quantified across three independent experiments. (C) Representative images of DAPI (blue) stained HFK micronuclei with or without γH2AX (red). The graph depicts mean ± standard error of the mean, * = *p*≤0.05 (Student’s *t*-test). In Δ8 E6, residues 132 to 136 were deleted. Micronuclei is abbreviated as MN.

Supplementary Figure 5:

Representative images iHFK micronuclei (A) with or without acetylated H3K27 (H3K27ac; red) staining and (B) with or without BrdU staining in iHFK cells grouped by micronuclear envelop morphology as determined by Lamin B1 (green) staining. DNA was stained with DAPI (white). (C) Representative images of DAPI (white) stained iHFK micronuclei with or without 53bp1 (red) in cells with indicated micronuclear membrane morphologies as determined by Lamin B1 (green) staining.

Supplementary Figure 6:

Chromothripsis plots that represent the four clusters from the chromothripsis analysis: High-confidence, Low-confidence, Abundant SVs, and Minimal Instability. All High-confidence chromosomes are shown A) 1, B) 2, C) 7, D) 11, and E) 13. Additionally, one chromosome from each other group is displayed as a representative. Chromosomes F) 3 (Low-confidence), G) 4 (Abundant SVs) and H) 14 (Minimal Instability). Vertical hashed bars show deletion-like events (orange), head-to-head inversions (black), and tail-to-tail inversions (green). Copy number (CN) variations are indicated by solid black bars ranging from 1-26 as indicated on the y-axis.

Supplementary Movie 1

This movie shows a representative iHFK LXSN cell expressing H2B-mCherry that undergoes mitosis with an anaphase bridge lasting 22.5 min. Images were captured every 7.5 minutes.

Supplementary Movie 2

This movie shows a representative iHFK 8 E6 cell (center-right at the start) expressing H2B-mCherry that undergoes mitosis with an anaphase bridge lasting 82.5 min that results in micronuclei formation. Images were captured every 7.5 minutes.

Supplementary Movie 3

This movie shows a representative iHFK LXSN cell (on the right) expressing RFP-NLS whose micronucleus loses RFP-NLS expression. Images were captured every 7.5 minutes.

Supplementary Movie 4

This movie shows an iHFK 8 E6 cell (bottommost cell) expressing RFP-NLS whose 2 micronuclei do not lose RFP-NLS expression. Images were captured every 7.5 minutes.

Supplementary Movie 5

This movie shows a representative iHFK LXSN cell expressing H2B-mCherry with a micronucleus that does not enter mitosis. Images were captured every 7.5 minutes.

Supplementary Movie 6

This movie shows a representative iHFK 8 E6 cell expressing H2B-mCherry with a micronucleus that enters and completes mitosis. Images were captured every 7.5 minutes.

## Acknowledgements

We appreciate KSU-CVM Confocal Core especially Joel Sanneman, for assisting with our immunofluorescence imaging, the Center on Emerging and Zoonotic Infectious Diseases for providing the live cell microscope, Michael Underbrink for providing hTERT-immortalized HFK cells, Emily Hatch for gifting the RFP-NLS plasmid.

Research reported in this publication was supported by the National Institute of General Medical Sciences (NIGMS) of the National Institutes of Health under award number P20 GM130448 and P20 GM103418, the Department of Defense CMDRP PRCRP CA160224, and made possible through generous support from the Johnson Cancer Research Center at Kansas State University. The content is solely the responsibility of the authors and does not necessarily represent the official views of the National Institutes of Health

